# Dopamine receptor type 2-expressing medium spiny neurons in the ventral lateral striatum have a non-REM sleep-induce function

**DOI:** 10.1101/2023.04.26.538415

**Authors:** Tomonobu Kato, Kenji F. Tanaka, Akiyo Natsubori

## Abstract

Dopamine receptor type 2-expressing medium spiny neurons (D2-MSNs) in the medial part of the ventral striatum (VS) induce non-REM (NREM) sleep from the wake state in animals. However, it is unclear whether D2-MSNs in the lateral part of the VS (VLS), which is anatomically and functionally different from the medial part of the VS, contribute to sleep-wake regulation. This study aims to clarify whether and how D2- MSNs in the VLS are involved in sleep-wake regulation. Our study found that specifically removing D2-MSNs in the VLS led to an increase in wakefulness time in mice during the dark phase using a diphtheria toxin-mediated cell ablation/dysfunction technique. D2-MSN ablation throughout the VS increased dark phase wakefulness time. These findings suggest that VLS D2-MSNs may induce sleep during the dark phase with the medial part of the VS. Next, our fiber photometric recordings revealed that the population intracellular calcium (Ca^2+^) signal in the VLS D2-MSNs increased during the transition from wake to NREM sleep. The mean Ca^2+^ signal level of VLS D2-MSNs was higher during NREM and REM sleep than during the wake state, supporting their sleep-inducing role. Finally, optogenetic activation of the VLS D2-MSNs during the wake state always induced NREM sleep, demonstrating the causality of VLS D2-MSNs activity with sleep-induction. Additionally, activation of the VLS D1-MSNs, counterparts of D2-MSNs, always induced wake from NREM sleep, indicating a wake- promoting role. In conclusion, VLS D2-MSNs could have an NREM sleep-inducing function in coordination with those in the medial VS.

**Significant statement:** The sleep-inducing function of D2-MSNs in the medial part of the ventral striatum (VS) has been previously reported; however, their function in the lateral part of the VS (VLS) has not been elucidated. We demonstrated that the diphtheria toxin-induced ablation of D2-MSNs in the VLS, as well as in the entire VS, increased wakefulness time in mice during the dark phase. VLS D2-MSNs had higher average Ca^2+^ signals during NREM and REM sleep than wake state via fiber photometric recording. Furthermore, optogenetic activation of VLS D2-MSNs during wake state induced NREM sleep in mice. In conclusion, D2-MSNs in the VLS have an NREM sleep-inducing function in coordination with those in the medial VS.

## Introduction

The regulation of sleep-wake behavior involves various neural populations, including the medium spiny neurons that express dopamine receptor type 2 (D2-MSNs) in the striatum (Oishi and Lazarus, 2017). D2-MSNs express receptors for adenosine, one of the sleep-promoting substances (Huang et al., 2014). These neurons are uniformly distributed throughout the striatum (Kemp JM and Powell TP, 1971); however, its functional manifestation is divided between the dorsal and ventral striatum (DS and VS), and further varies depending on the subregion within the DS/VS, such as during reward processing (Cox and Witten, 2019). Thus, the sleep-wake regulatory function of these neurons could also be exhibited differently in each striatal subregion.

In the DS, a region primarily involved in motor control in animals, D2-MSNs contribute to non-REM (NREM) sleep induction in the rostral, centromedial, and centrolateral subregions, but not in the caudal subregion, and their sleep-inducing role is exhibited only in the dark phase (Yuan et al., 2017). In contrast, in the VS, it is not clear in all subregions whether the D2-MSNs are involved in sleep-wake control. The VS can be anatomically and functionally divided into at least three subregions: the nucleus accumbens (NAc) medial shell, referred to as the ventral medial striatum (VMS), NAc core, and ventral lateral striatum (VLS; also known as NAc lateral shell). Previous studies have reported that D2-MSNs in the NAc core subregion contribute to NREM sleep induction in mice (Oishi et al., 2017; Luo et al., 2018). Another study has suggested that these neurons in the NAc medial shell have the same function (Satoh et al., 1999), and mediates caffeine-induced arousal via the adenosine A_2A_ receptor which co-express with dopamine D2 receptor (Lazarus et al., 2011). However, it is unclear whether D2-MSNs in the VLS subregion play a role in regulating sleep-wake cycles in animals. The VLS D2-MSNs receive distinctive input from cortical glutamatergic neurons in the insular cortex and dopaminergic neurons in the ventral tegmental area (VTA) (Hunnicutt et al., 2016; Mingote et al., 2019), and specifically constitute neuronal circuitry with parvalbumin-expressing interneurons (Yoshida et al., 2020). This could result in distinct firing patterns and functions in reward processing compared to those in the medial part of the VS (Tsutsui-Kimura et al., 2017; Yang et al., 2018; Chen et al., 2021). This anatomical and functional uniqueness of the VLS predicts a distinct role for VLS D2-MSNs in sleep-wake control.

To elucidate the role of VLS D2-MSNs in sleep-wake regulation, we evaluated the sleep-wake architecture using a 24-hour polysomnogram in mice, in which the D2-MSNs were spatiotemporally progressively ablated from the specific VLS to the entire VS based on DOX-dependent diphtheria toxin induction in these neurons (D2-DTA: Tsutsui-Kimura et al., 2017). Next, we investigated the Ca^2+^ signal patterns of the D2- MSNs in the VLS during physiological sleep-wake states in mice. Furthermore, we conducted optogenetic activation of the VLS D2-MSNs to reveal a causal relationship between the activity of these neurons and sleep-wake regulation. Our findings provide evidence that D2-MSNs in the VLS are involved in sleep induction in animals, in cooperation with those in the medial part of the VS.

## Materials and Methods

### Ethics statement

All animal procedures were conducted following the National Institutes of Health Guide for the Care and Use of Laboratory Animals and were approved by the Keio University Animal Experiment Committee in compliance with the Keio University Institutional Animal Care and Use Committee (approval numbers: A2022-315).

### Animals

Experiments were conducted with 8–14-month-old male and female mice. All mice were maintained on a 12:12 h light/dark cycle (lights on at 08:00; luminous flux, 120 lm). Polysomnographic recordings were performed on all days (start time at 08:00 [ZT0]), fiber photometric recordings were performed during the light phase (ZT0-8), and optogenetic manipulation was performed during the light and dark phase (ZT0-8 and ZT12-16). *Drd2*-DTA mice (*Drd2*-tTA::tetO-DTA [diphtheria toxin A]; hereafter referred to as D2-DTA) were obtained by crossing *Drd2*-tTA mice (Tsutsui-Kimura et al., 2017) and tetO-DTA mice (Lee et al., 1998). D2-DTA mice were fed doxycycline (DOX)-containing chow until the start of the experiment. Upon switching to normal chow (DOX-off day 0), tTA-mediated DTA induction was initiated as previously described (Tsutsui-Kimura et al., 2017). D2-YC mice (*Drd2*-tTA::tetO-YCnano50 double-transgenic mice) were generated by crossing *Drd2*-tTA and tetO-YCnano50 mice (Kanemaru et al., 2014). D2-ChR2 mice (*Drd2*-tTA::tetO-ChR2(C128S)-EYFP double transgenic mice) were generated by crossing *Drd2*-tTA and tetO-ChR2 mice (Tanaka et al., 2012). D1-ChR2 mice (*Pde10a2*-tTA::tetO- ChR2(C128S)-EYFP; *Adora2a*-Cre triple-transgenic mice) were obtained by crossing *Pde10a2*-tTA mice, tetO- ChR2(C128S)-EYFP mice, and *Adora2a*-Cre mice. The genetic backgrounds of all transgenic mice were mixed C57BL6 and 129 SvEvTac.

### Surgical procedure

Surgeries were performed using a stereotaxic apparatus (SM-6M-HT and SM-15R/L, Narishige). Mice were anesthetized with a mixture of ketamine and xylazine (100[and 10[mg/kg, respectively). Body temperature during surgery was maintained at 37[±[0.5[°C using a heating pad (FHC-MO, Muromachi Kikai). Mice received permanent electroencephalography (EEG) and electromyography (EMG) electrodes for polysomnography. Three pits were drilled into the skull using a carbide cutter (drill diameter: 0.8 mm). We implanted two electrodes in each subject, each consisting of a 1.0-mm diameter stainless steel screw that served as an EEG electrode. One implant was placed over the right frontal cortical area (AP: +1.0 mm; ML: +1.5 mm) as a reference electrode, while the other was placed over the right parietal area (AP: +1.0 mm anterior to lambda; ML: +1.5 mm) as a signal electrode. Another electrode was placed over the right cerebellar cortex (AP: −1.0 mm posterior to lambda; ML: +1.5 mm) as the ground electrode. Two silver wires (AS633; Cooner Wire Company, USA) were placed bilaterally into the trapezius muscles and served as the EMG electrodes. An optic fiber was inserted into the VLS at the following coordinates relative to bregma: AP, +1.1 mm; ML, +1.8 mm; DV, +4.4 mm from the skull unilaterally for photometry, AP, +1.1 mm; ML, ±1.9 mm; DV, +3.1 mm from the brain surface bilaterally for optogenetic manipulation. Finally, the electrode and optical fiber cannula assembly were anchored and fixed to the skull using SuperBond (Sun Medical Co.).

### EEG/EMG recordings

The EEG/EMG signals were amplified (gain ×1000) and filtered (EEG:1-300 Hz, EMG:10-300 Hz) using a DC/AC differential amplifier (AM-3000, AM systems). The input was received via an input module (NI-9215, National Instruments), digitized at a sampling rate of 1000 Hz using a data acquisition module (cDAQ-9174, National Instruments), and recorded using a custom-made LabVIEW program (National Instruments). We habituated the 24-hour EEG/EMG recordings more than three times, and REM sleep (see the vigilance state assessment) was often observed when we started the experiment.

### Fiber photometry

The ratiometric fiber photometry method has been described previously (Natsubori et al., 2017; Kato et al., 2022). An excitation light (435 nm; silver light-emitting diode, Prizmatix) was reflected off a dichroic mirror (DM455CFP, Olympus), focused with a 2× objective lens (numerical aperture 0.39, Olympus) and coupled into an optical fiber (400 mm diameter, 0.39 numerical aperture; catalog #M79L01, Thorlabs) through a pinhole (diameter 400 μm). The light-emitting diode power was <200 μW at the fiber tip. The cyan and yellow fluorescence emitted by the YC-nano50 was collected via an optical fiber cannula, divided by a dichroic mirror (DM515YFP, Olympus) into cyan (483/32 nm band path filters, Semrock) and yellow (542/27 nm), and detected using a photomultiplier tube (H10722-210, Hamamatsu Photonics). The fluorescence signals were digitized using a data acquisition module (cDAQ-9174, National Instruments) and recorded simultaneously using a custom-made LabVIEW program (National Instruments). Signals were collected at a sampling frequency of 1000 Hz.

### Optogenetic manipulation

An optical fiber (numerical aperture 0.39, Thorlabs) was inserted through the guide cannula. Blue (470 nm) and yellow (575 nm) light were generated using a SPECTRA 2- LCR-XA light engine (Lumencor). The blue and yellow light power intensities at the tip of the optical fiber were 1–2 and 3–4 mW, respectively. Using EEG and EMG monitoring, we illuminated one second of blue light to open the step-function-type opsin ChR2(C128S) (Berndt et al., 2009) of D2- or D1-ChR2(C128S) mice during the wake or NREM state lasting more than 10 s respectively. In control trials, yellow light was used instead of blue light.

### Mouse vigilance state assessment

EEG/EMG signals were analyzed using MATLAB (MathWorks, MA, USA). Power spectral EEG data were obtained using multitaper spectral estimation (McCoy et al., 1998). A power spectral profile over a 1-50–Hz window was used for the analysis. We detected each sleep-wake state scored offline by visually characterizing 10-s epochs for 24-hour polysomnography of D2-DTA mice and 1-s epochs for fiber photometric recording of D2-YC mice and optogenetic experiments with D2- and D1-ChR2 mice (Funato et al., 2016; Kato et al., 2022). The state determination criteria were as follows: the wake state was characterized by a low-amplitude fast EEG and high-amplitude variable EMG. NREM sleep is characterized by a high-amplitude delta (1-4 Hz) frequency EEG and low-amplitude tonus EMG. REM sleep was staged using theta (6-9 Hz) dominant EEG and electromyography (EMG) atonia (Tobler et al., 1997).

### Colorimetric In situ hybridization (ISH)

Mice were deeply anesthetized with ketamine (100 mg/kg) and xylazine (10 mg/kg) and perfused with a 4% paraformaldehyde phosphate buffer solution. After removing the brains from the skull, they were fixed overnight in the same solution. The brains were then cryoprotected in 20% sucrose overnight, frozen, and sectioned to 25-μm thickness on a cryostat. The sections were mounted on silane-coated glass slides (Matsunami Glass), and then treated with proteinase K (40 μg[ml^−1^; Merck). After washing and acetylation, sections were incubated with digoxigenin (DIG)-labeled complementary RNA (cRNA) probes. After the sections were washed in buffers with serial differences in stringency, they were incubated with an alkaline phosphatase- conjugated anti-DIG antibody (11093274910, 1:5,000; Roche). The cRNA probes were visualized using freshly prepared colorimetric substrate (NBT/BCIP; Roche). Nuclear Fast Red (Vector Laboratories) was used for counterstaining (Tanaka et al., 2012). Probes for *DTA*, *Drd1*, and *Drd2* were used in this study. Images were obtained using an all-in-one microscope (BZ-X710; Keyence).

### Data processing

All animals and trials were randomly assigned to experimental conditions. The experimenters were not blinded to the experimental conditions during data collection and analysis. Mice were excluded when the optical fiber position was incorrectly targeted. We used custom-made programs in MATLAB for signal processing.

### EEG/EMG signal analysis

The EEG signals were bandpass (FIR)–filtered (Kaiser window). The EEG frequency bands were set as follows: delta: 1-4 Hz; theta: 6-9 Hz; sigma: 9-12 Hz; beta: 12-30 Hz; and gamma: 30-50 Hz (Choi et al., 2010). The power of each EEG frequency band was obtained using the square of the amplitude.

### Fluorescence signal analysis

For fiber photometric signal analysis, the yellow and cyan fluorescence signals were detrended. Further, we used the YC ratio (*R*), which is the ratio of yellow to cyan fluorescence intensity, calculated as the population Ca^2+^ signals, and then Z-scored. For the power spectral analysis, fiber photometric signals (YC signals) were extracted during the uninterrupted wake, NREM sleep, and REM sleep bouts that lasted >100 s, high pass filtered with a 0.01 Hz cutoff frequency, and calculated with a wavelet transform using a Morse wavelet from 0.01 to 1 Hz with time-bandwidth parameter (*P^2^* = 10). The peak frequency is determined as the local maximum of the power spectrum. For the state transition analysis, YC signals were extracted during uninterrupted wake, NREM sleep, and REM sleep bouts that lasted >20 s, and normalized using the mean and standard deviation of YC signals from 20 s before the transition to the transition point.

### Statistical analysis

All experimental data were analyzed using the following parametric statistics: Student’s t-test (independent samples t-test), paired t-test, paired t-test with Bonferroni correction, and repeated measures ANOVA followed by the Tukey-Kramer post-hoc test.

### Data availability

The datasets generated and/or analyzed in the current study are available from the corresponding author upon reasonable request.

## Results

### Ablation of D2-MSNs in the VLS causes an increase in wakefulness time in the dark phase

To investigate the role of D2-MSNs in the sleep-wake regulation within the ventral lateral striatum (VLS), we conducted repeated 24-hour polysomnography on *Drd2*- tTA::tetO-DTA mice (D2-DTA; Tsutsui-Kimura et al., 2017). In this mouse line, tTA- mediated diphtheria toxin expression is induced in D2-MSNs, which is initiated specifically in the VLS and follows spatial progress within the ventral striatum (VS) in doxycycline (DOX)-dependent manner (**Fig. 1a**). Upon switching to normal chow from DOX-containing chow (DOX-off day 0), DTA induction is first observed only in the VLS (corresponding to the NAc lateral shell), and the DTA induction area gradually expands toward the medial part of the VS (corresponding to the NAc medial shell and core) with each passing day. Based on the histological and behavioral assessments of the D2-DTA mice in a previous study (Tsutsui-Kimura et al., 2017), the VLS D2-MSNs could be regarded as a hypofunction condition on DOX-off day 5 and as a dysfunction condition (cell death) on DOX-off day 7, respectively (**Fig. 1b**). On DOX-off day 10, the D2-MSNs in the entire VS could fall into a hypofunctional condition with VLS D2- MSNs dysfunction. We histologically confirmed the *DTA* mRNA signal expressed in the entire VS on DOX-off day 10, indicating the hypofunctional condition of the D2-MSNs in the entire VS (**Fig. 1d**). The VLS-specific loss of *Drd2* mRNA signal indicated cell death of the VLS D2-MSNs (**Figs. 1c, d**).

**Figure 1.**
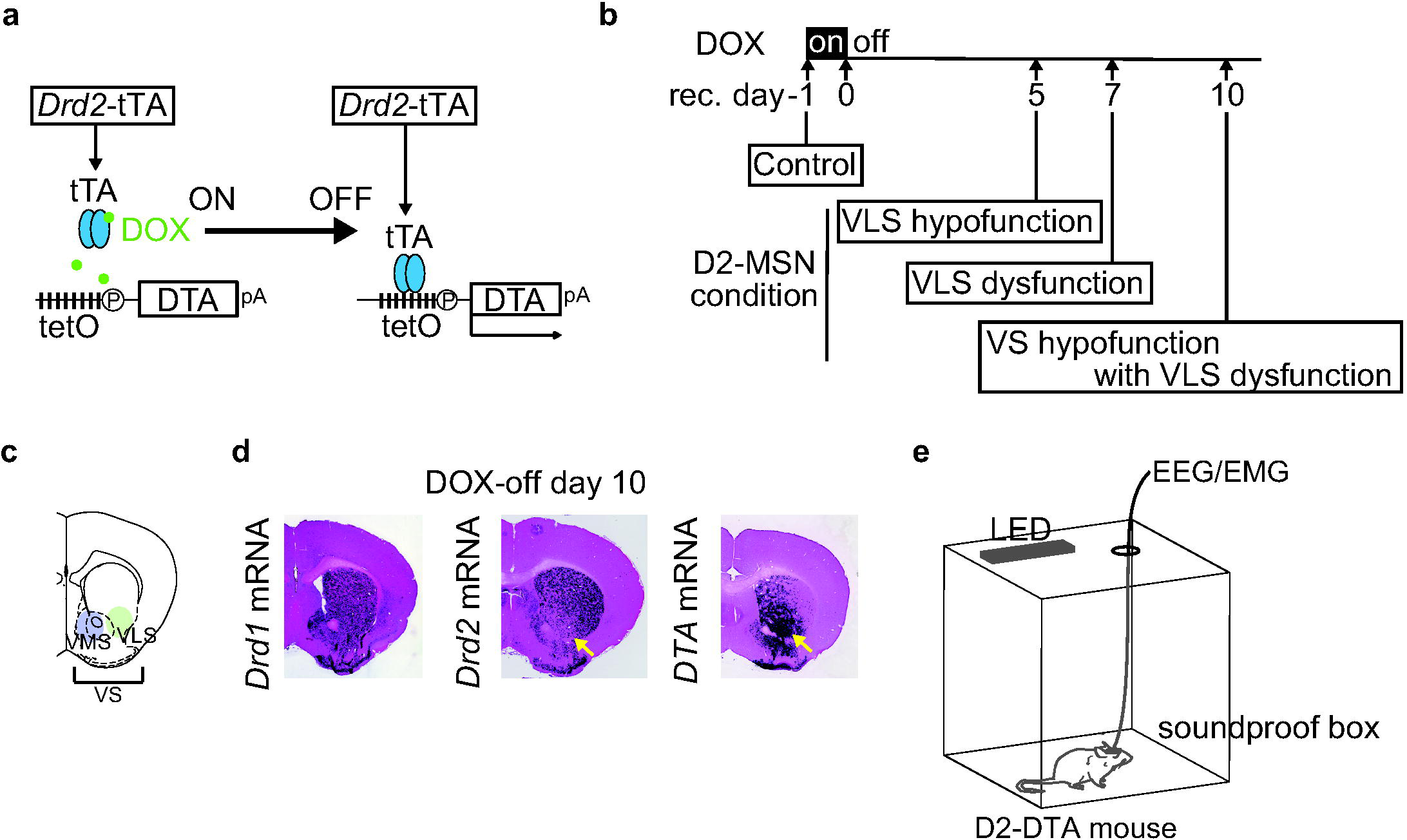
Spatiotemporally specific the VS D2-MSNs ablation. (a) DOX-controllable DTA expression. *Drd2*-tTA::tetO-DTA (D2-DTA) mice were fed with doxycycline (DOX)-containing chow until the start of the experiment (DOX-ON). *DTA* mRNA expression started when DOX-chow was replaced with normal chow (DOX-OFF). (b) Time course of the VS D2-MSNs ablation, and timing of the polysomnographic recordings. (c) Schematic illustration of the VS, including the VMS and the VLS. (d) *Drd1, Drd2, and DTA* mRNA levels in the striatum on DOX-off day 10. Purple denotes the mRNA signal. Left: *Drd1* mRNA expression throughout the striatum. Middle: *Drd2* mRNA expression in the VLS disappears (yellow arrow). Right: *DTA* mRNA expression was observed throughout the VS (yellow arrow). (e) Experimental setup. We conducted 24-hour EEG/EMG recording in a soundproof box and set a 12:12 h light/dark cycle using LED light.

We performed 24-hour polysomnography recordings to evaluate the effect of progressive ablation of the VS D2-MSNs in the D2-DTA mice on the sleep-wake architecture (**Fig. 1e**). The recordings were done on DOX-off day -1 (control), DOX-off days 5 and 7 (functional ablation of VLS D2-MSNs), and DOX-off day 10 (ablation of D2-MSNs in the entire VS) to assess the impact of these ablations (**Fig. 1b**). We observed that the daily amount of wakefulness time significantly increased on DOX-off day10, alongside a decrease in NREM sleep time, compared to those on DOX-off day - 1 (\**p* < 0.05, Repeated Measures ANOVA followed by the Tukey–Kramer *post-hoc* test; **Fig. 2a**). However, the daily sleep-wake time did not significantly change on DOX-off days 5 and 7. Next, we separately analyzed the effect of DOX-dependent D2- MSNs ablation on sleep-wake architecture in the light and dark phases. In the light phase, there were no significant differences in the percentage of time spent in the wake, in NREM sleep, or in REM sleep (**Fig. 2b**). During the dark phase, we observed a significant increase in the percentage of time spent in the wake on DOX-off day 5, which continued to increase up to DOX-off day 10, along with decreases in both NREM and REM sleep (\**p* < 0.05, Repeated Measures ANOVA followed by the Tukey– Kramer *post-hoc* test; **Fig. 2b**). This phenotype of increased wake time and decreased sleep time due to D2-MSNs ablation was most pronounced in the first half of the dark phase (\**p* < 0.05, paired *t*-test at each time point with Bonferroni correction; **Fig. 2c**).

**Figure 2.**
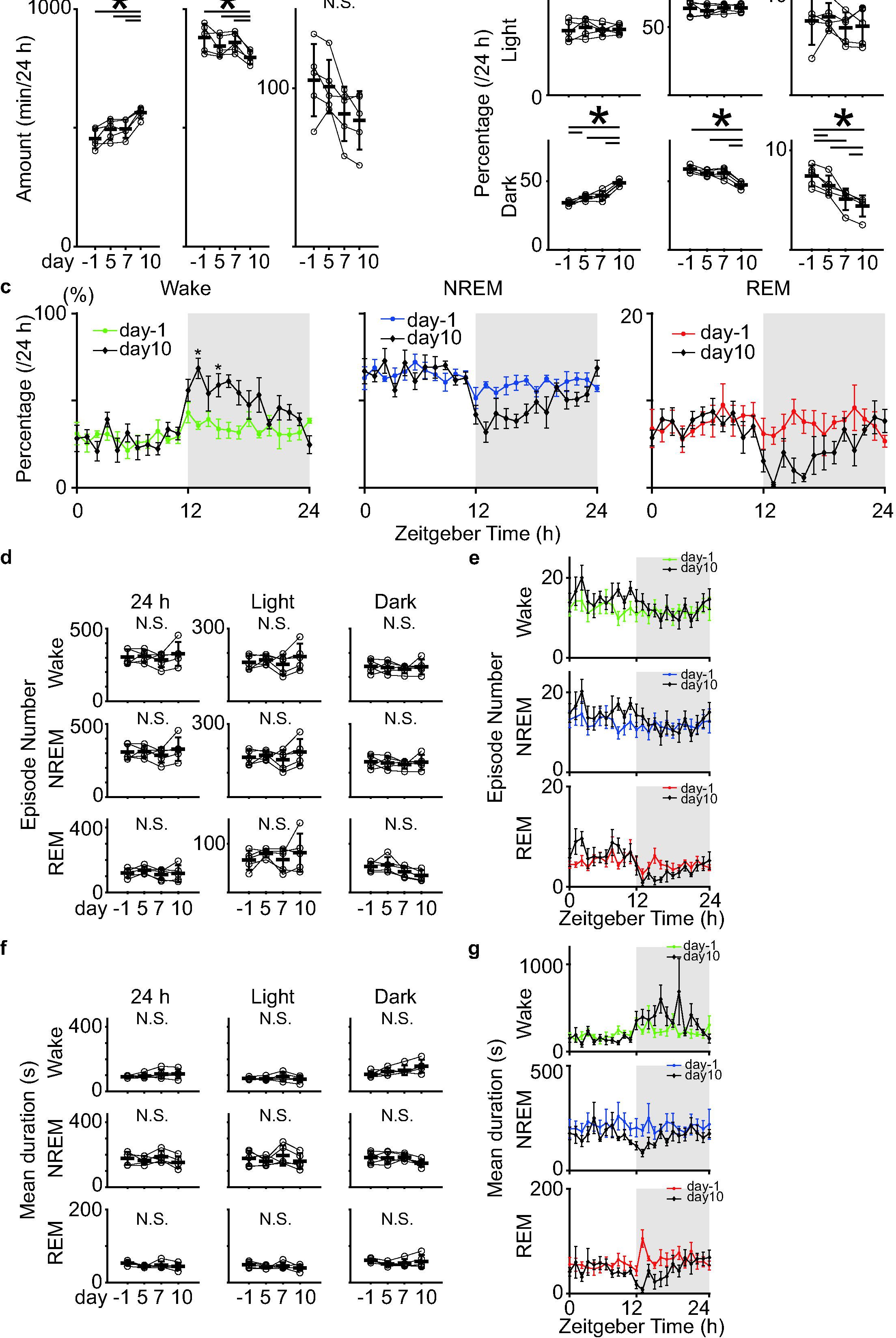
Analysis of sleep structure before and after the VLS/VS D2-MSNs ablation. _(a)_ Total amount of wake, NREM sleep, and REM sleep time in control (DOX-off day - 1) and DOX-off days 5, 7, and 10 (n = 5 mice). Repeated-measures ANOVA was followed by the Tukey–Kramer *post-hoc* test. Wake: *p*_day-1 vs day5_ = 0.17, *p*_day-1 vs day7_ = 0.29, *p*_day-1 vs day10_ = 6.0×10^-3^, *p*_day5 vs day7_ = 0.99, *p*_day5 vs day10_ = 8.0×10^-3^, *p*_day7 vs day10_ = 8.2×10^-3^; NREM: *p*_day-1 vs day5_ = 0.33, *p*_day-1 vs day7_ = 0. 83, *p*_day-1 vs day10_ = 0.03, *p*_day5 vs day7_ =0.47, *p*_day5 vs day10_ = 0.03, *p*_day7 vs day10_ = 7.0×10^-3^; REM: *p*_day-1 vs day5_ = 0.86, *p*_day-1 vs day7_ = 0.14, *p*_day-1 vs day10_ = 0.16, *p*_day5 vs day7_ = 0.14, *p*_day5 vs day10_ = 0.20, *p*_day7 vs day10_ = 0.60. (b) Time spent in each sleep state during the light and dark phases in control (DOX-off day -1) and DOX-off mice on days 5, 7, and 10 (n = 5 mice). Repeated Measures ANOVA, Wake (light): *p* = 0.72; NREM (light): *p* = 0.43; REM (light): *p* = 0.41. Repeated-measures ANOVA was followed by the Tukey–Kramer *post-hoc* test. Wake (dark): *p*_day-1 vs day5_ = 4.5×10^-3^, *p*_day-1 vs day7_ = 0.14, *p*_day-1 vs day10_ = 1.1×10^-3^, *p*_day5 vs day7_ = 0.70, *p*_day5 vs day10_ = 3.6×10^-4^, *p*_day7 vs day10_ = 5.1×10^-4^; NREM (dark): *p*_day-1 vs day5_ = 0.13, *p*_day-1 vs day7_ = 0.53, *p*_day-1 vs day10_ = 6.4×10^-3^, *p*_day5 vs day7_ = 1, *p*_day5 vs day10_ = 3.4×10^-3^, *p*_day7 vs day10_ = 3.9×10^-4^; REM (dark): *p*_day-1 vs day5_ = 0.03, *p*_day-1 vs day7_ = 8.3×10^-3^, *p*_day-1 vs day10_ = 6.7×10^-3^, *p*_day5 vs day7_ = 0.12, *p*_day5 vs day10_ = 4.7×10^-2^, *p*_day7 vs day10_ = 0.01. (c) Daily variations in wake time (left), NREM sleep (middle), REM sleep (right) every hour in control (DOX-off day -1), and DOX-off day 10 (n = 5 mice). Paired *t*-test at each time point with Bonferroni correction for family wise error rate of *p* = 0.05. (d) Episode numbers of wake, NREM sleep, and REM sleep for 24 h, light, and dark phases in control (DOX-off day -1) and DOX-off days 5, 7, and 10 (n = 5 mice). Repeated-measures ANOVA. Wake (24 h): *p* = 0.62; wake (light), *p* = 0.36; wake (dark), *p* = 0.89; NREM (24 h): *p* = 0.60; NREM (light), *p* = 0.33; NREM (dark), *p* = 0.88; REM (24 h): *p* = 0.30; REM (light), *p* = 0.42; REM (dark), *p* = 0.88. (e) Daily variations in number of episodes during each sleep/wake state in control (DOX-off day -1), DOX-off day 10 (n = 5 mice; Paired *t*-test at each time point with Bonferroni correction for family wise error rate of *p* = 0.05). (f) Mean duration of each bout of wake, NREM sleep, and REM sleep for 24 h, light, and dark phases in control (DOX-off day -1) and DOX-off days 5, 7, and 10 (n = 5 mice). Repeated-measures ANOVA. Wake (24 h): *p* = 0.21, Wake (light): *p* = 0.22, Wake (dark): *p* = 0.04 [no significant difference between each group using the Tukey–Kramer *post-hoc* test], NREM (24 h): *p* = 0.20, NREM (light): *p* = 0.29, NREM (dark): *p* = 0.16, REM (24 h): *p* = 0.14, REM (light): *p* = 0.37, REM (dark): *p* = 0.16. (g) Daily variations in mean duration during each sleep/wake state in control (DOX-off day -1) and DOX-off day 10 (n = 5 mice). Paired *t*-test at each time point with Bonferroni correction for family wise error rate of *p* = 0.05. Error bars indicate SEM; * *p* < 0.05.

To determine the characteristics of sleep-wake alteration induced by D2-MSNs ablation, we calculated the number of episodes of wake, NREM sleep, and REM sleep, but they were not affected (*p < 0.05, Repeated Measures ANOVA followed by the Tukey–Kramer *post-hoc* test; **Fig. 2d**, paired *t*-test at each time point with Bonferroni correction; **Fig. 2e**). The mean duration of the sleep/wake bout showed a gradual increasing trend in the dark phase under progressive D2-MSNs ablation, although this was not statistically significant (\**p* < 0.05, Repeated Measures ANOVA followed by the Tukey–Kramer *post-hoc* test; **Fig. 2f**, paired *t*-test at each time point with Bonferroni correction; **Fig. 2g**).

Next, we investigated the EEG changes in sleep or wakefulness that were altered by VS D2-MSN ablation. When we compared the EEG spectrum during each sleep- wake state throughout the day and in the light/dark phases between the control (DOX- off day -1) and DOX-off day 10, there were no significant differences in any of the power bands of the EEG signal (delta, theta, sigma, beta, and gamma) (\**p* < 0.05, paired *t*-test; delta and theta ranges are shown in **Fig. 3**).

**Figure 3.**
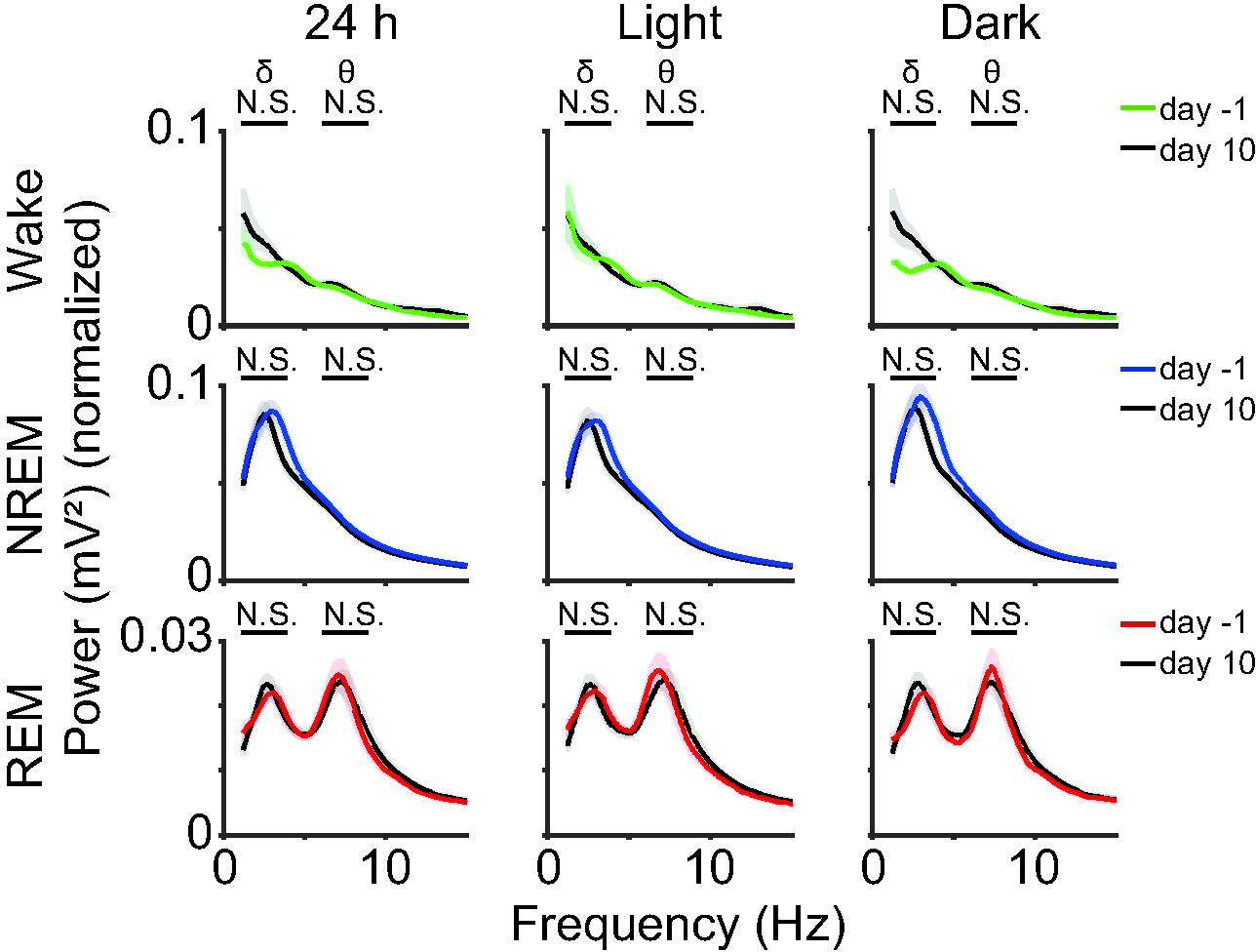
Comparison of EEG power spectrum before and after the VS D2-MSNs ablation. EEG spectra before (DOX-off day -1) and after (DOX-off day10) the VS D2-MSNs ablation during each sleep/wake state (n = 5 mice). Paired *t*-test. Wake, delta: *p*_24 h_ = 0.30, *p*_light_ = 0.75, *p*_dark_ = 0.32, theta: *p*_24 h_ = 0.12, *p*_light_ = 0.42, *p*_dark_ = 0.36; NREM, delta: *p*_24 h_ = 0.14, *p*_light_ = 0.20, *p*_dark_ = 0.16, theta: *p*_24 h_ = 0.16, *p*_light_ = 0.29, *p*_dark_ = 0.08; REM, delta: *p*_24 h_ = 0.96, *p*_light_ = 0.51, *p*_dark_ = 0.48, theta: *p*_24 h_ = 0.98, *p*_light_ = 0.98, *p*_dark_ = 0.98. Colored shades indicate SEM; * *p* < 0.05.

### VLS D2-MSNs displayed higher Ca^2+^ signal levels during NREM and REM sleep compared to wake

Our ablation study revealed that D2-MSNs in the VLS may be involved in sleep-wake regulation. Thus, we monitored the compound intracellular calcium (Ca^2+^) signal patterns of the VLS D2-MSNs to determine their activity patterns across sleep-wake states and during sleep/wake stage transitions in mice. We used a fiber photometry system to monitor intracellular Ca^2+^ signals from the VLS D2-MSNs in freely moving mice (**Figs. 4a, b**). We used transgenic mice expressing the FRET-based ratiometric Ca^2+^ indicator YC-nano50 (Horikawa et al., 2010), in D2-MSNs under the control of the *Drd2* promoter (*Drd2*-tTA::tetO-YCnano50: D2-YC mice; Natsubori et al., 2017). The ratio of yellow to cyan fluorescence intensity (YC ratio) represented the intracellular Ca2+ signal of D2-MSNs.

**Figure 4.**
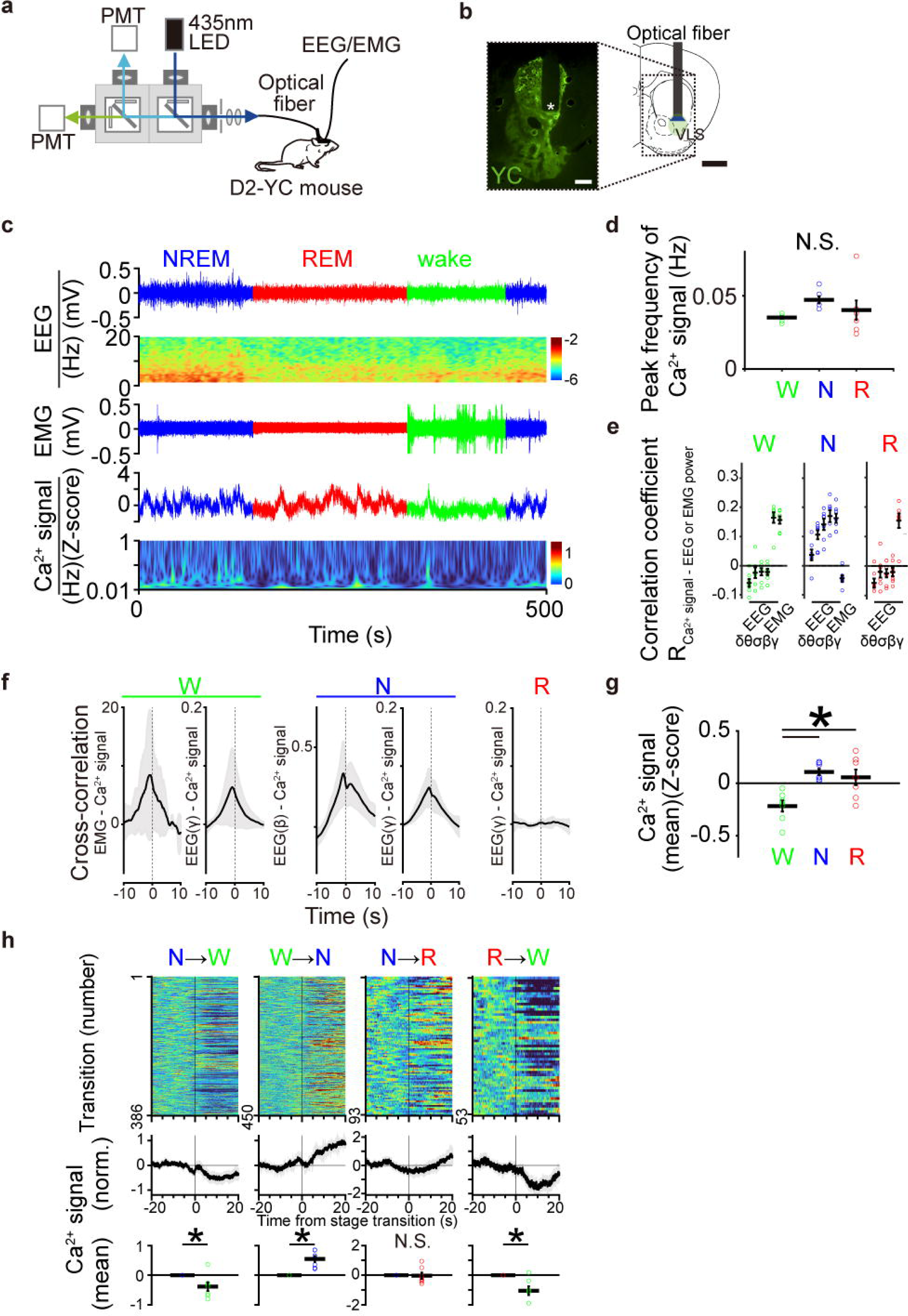
Population Ca^2+^ signals of the VLS D2-MSNs across the sleep–wake states. (a) Schematic illustration of the fiber photometry system for monitoring intracellular Ca^2+^ signals of VLS D2-MSNs in D2-YC mice. The EEG and EMG were recorded simultaneously. PMT, Photomultiplier tube. (b) YCnano50 fluorescent expression in D2-YC mice (left) and a schematic of the fiber photometric recording site (right) in the VLS. The asterisks indicate the tip of the optical fiber. Scale bar, 1 mm. (c) Representative examples of EEG signal trace and power spectrogram, EMG signal, and population Ca^2+^ signal trace and spectrogram of VLS D2-MSNs across the sleep–wake states. The green line shows wakefulness, the blue line shows NREM sleep and the red line shows REM sleep. (d) Peak frequency of Ca^2+^ signals in VLS D2-MSNs during each sleep/wake state (28 recordings from 7 mice; one-way ANOVA followed by Tukey–Kramer *post-hoc* test, wake vs. NREM, *p* = 0.13; NREM vs. REM, *p* = 0.47; wake vs. REM, *p* = 0.67). (e) Correlation coefficient between the Ca^2+^ signals and each frequency band (δ, θ, σ, β and γ) of EEG power or EMG power during wake and NREM sleep, each frequency band (δ, θ, σ, β and γ) of EEG power during REM sleep. (f) Cross-correlation between EMG power and Ca^2+^ signals during wake, between EEG gamma power and Ca^2+^ signals during wake, between EEG beta and gamma power and Ca^2+^ signals during NREM sleep, and between EEG gamma power and Ca^2+^ signals during REM sleep. (g) Mean Ca^2+^ signal levels of the VLS D2-MSNs across the sleep-wake states (28 recordings from 7 mice; one-way ANOVA followed by Tukey–Kramer *post-hoc* test, wake vs. NREM, *p* = 1.7×10^-3^; NREM vs. REM, *p* = 0.80; wake vs. REM, *p* = 7.0×10^-3^). (h) The Ca^2+^ signal of the VLS D2-MSNs aligned with each sleep/wake transition. Upper panel: Ca^2+^ signal fluctuations during individual transitions with color-coded fluorescence intensity (NREM to wake, n = 386; wake to NREM, n = 450; NREM to REM, n = 93; REM to wake, n = 53 events from 7 mice). Middle panel: average Ca^2+^ signals from all transitions. Lower panel, mean of Ca^2+^ signal levels before and after 20 s from each stage transition (VLS: *p*_NREM-wake_ = 3.7×10^-3^, *p*_wake-NREM_ = 0.03, *p*_NREM-REM_ = 0.33, *p*_REM-wake_ = 0.02. Paired *t*-test). Gray shading and error bars indicate SEM; \**p* < 0.05. W, wake; N, NREM sleep; R, REM sleep.

We found that the VLS D2-MSNs displayed infra-slow periodic Ca^2+^ signal fluctuations during wake, NREM sleep, and REM sleep (**Figs. 4c, d**). The frequency of these events did not show a significant variation between the different sleep/wake states but was slightly higher during NREM sleep compared to wakefulness and REM sleep (**Fig. 4d**). Next, we investigated the temporal relationship between Ca^2+^ signal fluctuations in the VLS D2-MSNs and EEG/EMG activity during each sleep/wake state. When we calculated the correlation coefficient between the Ca^2+^ signal and the EEG or EMG power, the Ca^2+^ signal in the VLS D2-MSNs shows positive correlation with EEG gamma power and EMG power during the wake (correlation coefficient: 0.16 ± 0.02 and 0.16 ± 0.01; mean ± S.E.M.), EEG beta and gamma power during the NREM sleep (correlation coefficient: 0.17 ± 0.02 and 0.16 ± 0.02; mean ± S.E.M.), and EEG gamma power during the REM sleep (0.15 ± 0.02; mean ± S.E.M.; **Fig. 4e**). Using cross-correlation analysis, we found that the Ca^2+^ signal in VLS D2-MSNs preceded the peak of EEG high-frequency or EMG power during both wake (lag time between peaks, EEG gamma power – Ca^2+^ signal: -0.83 ± 0.10 s, EMG power – Ca^2+^ signal: -0.82 ± 0.15 s; mean ± S.E.M.) and during the NREM sleep (lag time between peaks, EEG beta power - Ca^2+^ signal: -0.31 ± 0.46 s, EEG gamma power - Ca^2+^ signal: -0.77 ± 0.18 s; mean ± S.E.M.; **Fig. 4f**). However, no such relationship was observed between the Ca^2+^ signal and EEG gamma power during REM sleep, likely because gamma power fluctuations are minimal during this state (**Fig. 4f**).

Furthermore, we focused on the averaged Ca^2+^ signal levels during each sleep/wake state, which was significantly higher during NREM and REM sleep than during wakefulness (**Figs. 4c, g**). Thus, we evaluated changes in the average Ca^2+^ signal level during the state transition (**Fig. 4h**). During the transition from NREM sleep to wake, the Ca^2+^ signal began to decrease, and from wake to NREM sleep, the Ca^2+^ signal began to increase. The Ca^2+^ signal did not change during the transition from NREM to REM sleep, and it began to decrease during the transition from REM sleep to wake (**Fig. 4h**). These Ca^2+^ signal pattern variations in the VLS D2-MSNs, particularly during NREM sleep induction, support the sleep-inducing function of these neurons, as indicated by our ablation experiment.

### Optogenetic activation of the VLS D2-MSNs during wake causes NREM sleep

Our ablation study and Ca^2+^ signal recordings suggest that VLS D2-MSNs have an NREM sleep-inducing function. Therefore, to determine the causal relationship between the VLS D2-MSN activity and NREM sleep induction, we performed optogenetic activation of the VLS D2-MSNs. We used transgenic mice in which only the D2-MSNs expressed the step-function-type variant of ChR2 (D2-ChR2 (C128S)) and artificially activated their VLS D2-MSNs (**Fig. 5a**). In this variant, short-term light induction cause to initiate photocurrent, and it terminated approximately 100 s after light induction because the “tau off” of C128S mutant ion channel is approximately 106 ± 9 s (Berndt et al., 2009). The D2-ChR2 (C128S) mice received 1 s of blue light illumination to open the ChR2 (C128S) in the bilateral VLS D2-MSNs or 1 s of yellow light as a control (**Fig. 5a**).

**Figure 5.**
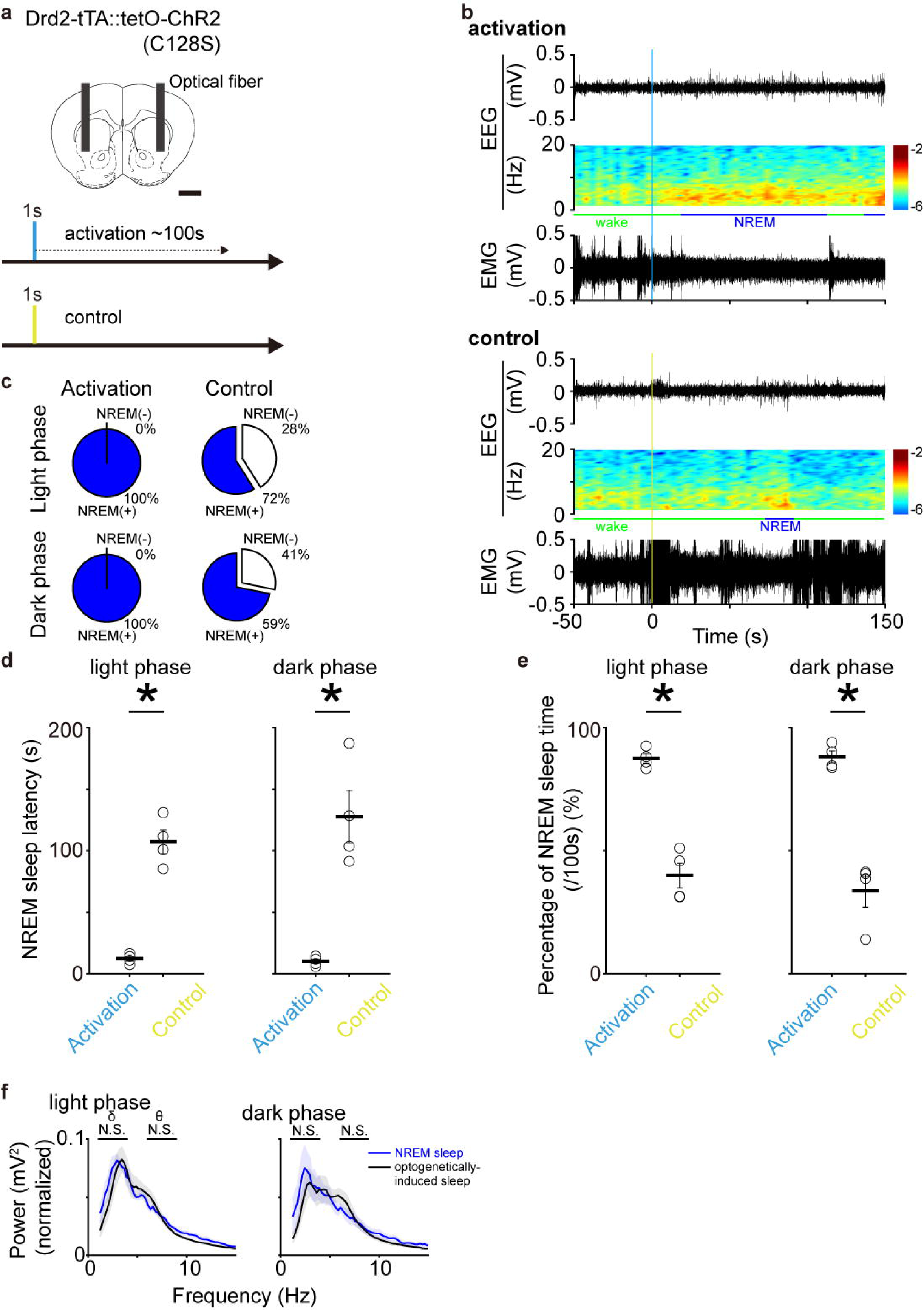
Sleep-wake state change by optogenetic activation of the VLS D2-MSNs. (a) Schematic illustration of optogenetic activation of VLS D2-MSNs in *Drd2-* ChR2(C128S) mice (hereafter referred to as D2-ChR2 mice). Scale bar, 1 mm. The blue and yellow shades indicate the illumination times. (b) Representative examples of EEG signal traces, power spectrograms, and EMG signals. Vertical blue and yellow shades indicate illumination times. The upper panel shows the optogenetic manipulation and the lower panel shows the control experiment. (c) NREM sleep induction rate which indicates whether NREM sleep occurred within 100 s after light illumination. Upper column shows a light phase and lower column shows a dark phase. In each column, left side shows the activation group (blue light illumination) and right side shows control group (yellow light illumination) (n = 4 mice, 5-10 manipulations/recordings, a total of 49 trials for activation [light phase], 51 trials for control [light phase], a total of 38 trials for activation [dark phase], 39 trials for control [dark phase]). (d) NREM sleep latency after the initiation of optogenetic activation of VLS D2-MSNs. The columns on the left and right show the light and dark phases, respectively. In each column, left bar shows the activation group, right bar shows the control group (n = 4 mice, 5-10 manipulations/recordings, a total of 49 trials for activation [light phase], 51 trials for control [light phase], a total of 38 trials for activation [dark phase], 39 trials for control [dark phase]; *p*_light_ = 6.9 × 10^-5^, *p*_dark_ = 1.5 × 10^-3^. independent *t*-test). (e) Percentage of NREM sleep time for 100 s after the initiation of VLS D2-MSNs photoactivation. The columns on the left and right show the light and dark phases, respectively. In each column, left bar shows the activation group, light bar shows the control group (n = 4 mice, 5-10 manipulations/recordings, a total of 49 trials for activation [light phase], 51 trials for control [light phase], a total of 38 trials for activation [dark phase], 39 trials for control [dark phase]; *p*_light_ = 1.2 × 10^-4^, *p*_dark_ = 2.4 × 10^-4^, independent *t*-test). (f) EEG spectrum of D2-ChR2 mice during natural NREM sleep (blue line; we extracted and averaged 200 s of NREM sleep before optogenetic activation) and optogenetically induced sleep (black line) during the light (left panel) and dark phases (right panel) (n = 4 mice, the same mice in Figs. 5c-e. delta: *p*_light_ = 0.63, *p*_dark_ = 0.94; theta: *p*_light_ = 0.59, *p*_dark_ = 0.75, independent *t*-test). Colored shades and error bars indicate SEM; * *p* < 0.05.

Optogenetic activation of VLS D2-MSNs during the wake state in mice always prompted changes in EEG/EMG activity and induced NREM sleep within 100 s after light illumination during both the light and dark phases; however, control light (yellow light) did not always induce these changes (**Figs. 5b, c**). The NREM sleep-induction latency by the VLS D2-MSNs photoactivation was 12.3 ± 2.0 and 10.2 ± 1.9 s in the light and the dark phase respectively, which was significantly shorter than those of the control light illumination (mean ± S.E.M, *p* = 6.9×10^-5^ and *p* = 1.5×10^-3^ [light and dark phase, respectively], independent *t*-test; **Fig. 5d**). The percentage of NREM sleep time during 100 s after the start of VLS D2-MSNs photoactivation were 87.6 ± 1.9 and 88.1 ± 2.4 % in the light and the dark phases, respectively, which were significantly higher than that of control light illumination (mean ± S.E.M, *p* = 1.2×10^-4^ and *p* = 2.4×10^-4^ [light and dark phase, respectively], independent *t*-test; **Fig. 5e**). Furthermore, EEG delta and theta powers were at the same level between natural NREM sleep and the optogenetically induced NREM sleep during both the light and dark phases (**Fig. 5f**). These results suggest that the VLS D2-MSNs activity induced NREM sleep in mice.

### Optogenetic activation of the VLS D1-MSNs induces wakefulness from NREM sleep

In the striatum, another population of MSNs, expressing dopamine receptor type 1 (D1-MSNs), is also distributed. While D2- and D1-MSNs in the dorsal striatum have opposing roles in motor control and reinforcement learning, in the VLS they have been reported to have cooperative roles in goal-directed behavior in mice (Kravitz et al 2010; Natsubori et al., 2017). Therefore, we tried optogenetic activation of the D1-MSNs in the VLS to compare the sleep-wake regulation functions of VLS D2- and D1-MSNs. We used D1-ChR2 mice (Pde10a2-tTA::tetO- ChR2(C128S)-EYFP; Adora2a-Cre triple-transgenic mice), which harboring ChR2(C128s) only in the D1-MSNs (Fig.6a). As a result, optogenetic activation of the VLS D1-MSNs during wake state did not cause any state changes (**Fig. 6b**). However, activation of the VLS D1-MSNs during NREM sleep always promptly induced wake in mice during the light and dark phases, whereas control light (yellow light) illumination did not always induce it (**Figs. 6c-f**). The wake-induction latency by the VLS D1-MSNs photoactivation was 0.15 ± 0.02 and 0 s in the light and the dark phase respectively (mean ± S.E.M, *p* = 0.01 and *p* = 0.03 [light and dark phase, respectively], independent *t*-test; **Fig. 6e**). The percentage of wake time during 100 s after the start of VLS D1-MSNs photoactivation were 96.7 ± 1.3 and 100 % in the light and the dark phases, respectively (mean ± S.E.M, *p* = 6.3 × 10^-6^ and *p* = 2.4 × 10^-5^ [light and dark phase, respectively], independent *t*-test; **Fig. 6f**). These results indicates that VLS D1-MSNs have a wake-inducing function and that VLS D2- and D1-MSNs encode opposing functions in sleep-wake regulation.

**Figure 6.**
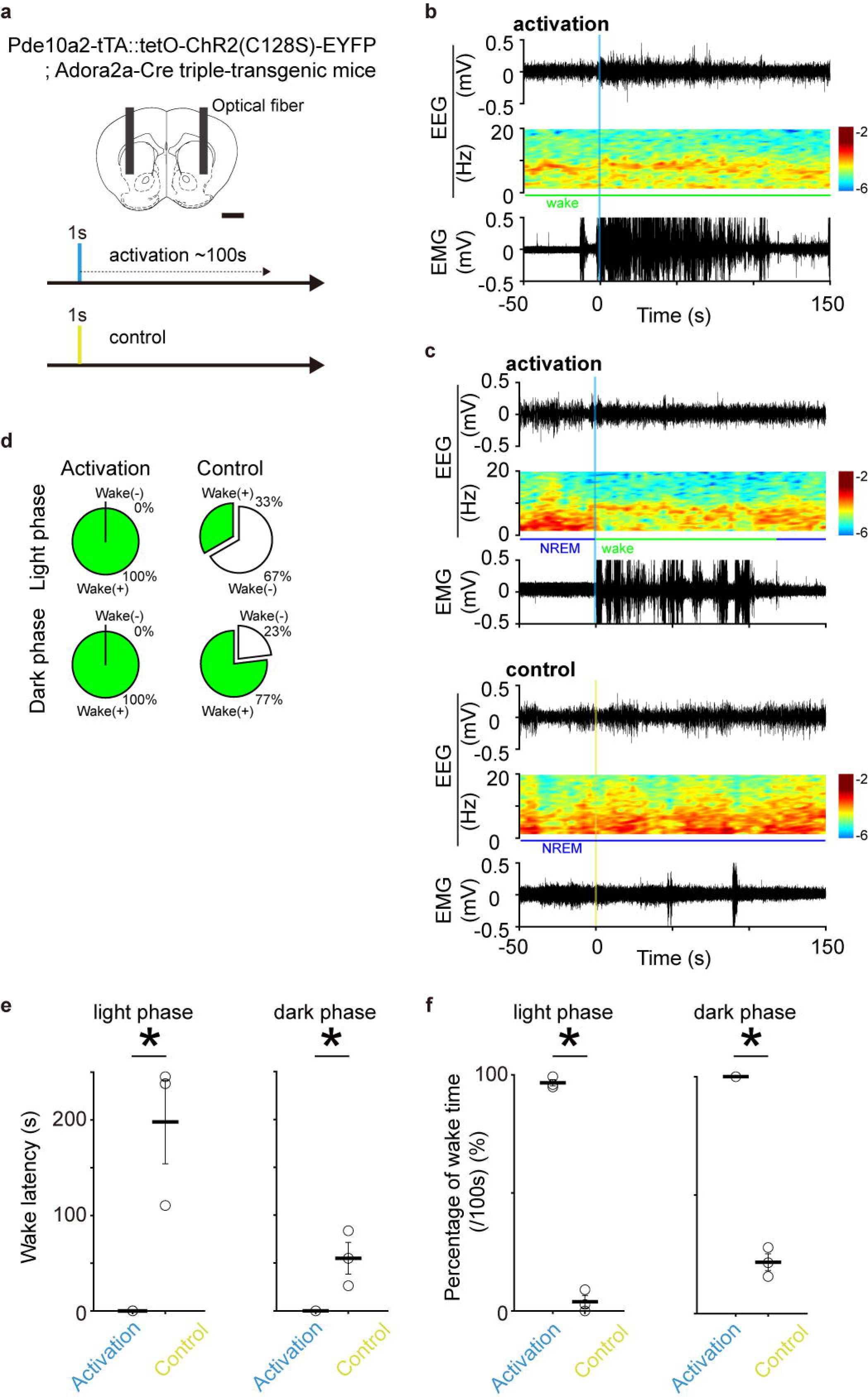
Sleep-wake state change by optogenetic activation of the VLS D1-MSNs. (a) Schematic illustration of optogenetic activation of VLS D1-MSNs in *Pde10a2*- tTA::tetO- ChR2(C128S)-EYFP; *Adora2a*-Cre triple-transgenic mice. Scale bar, 1 mm. The blue and yellow shades indicate the illumination times. (b) Representative examples of EEG signal traces, power spectrograms, and EMG signals before and after optogenetic manipulation during the wake. Vertical blue shade indicates illumination times. (c) Representative examples of EEG signal traces, power spectrograms, and EMG signals before and after optogenetic manipulation during NREM sleep. Vertical blue shade indicates illumination times. The upper panel shows the optogenetic manipulation and the lower panel shows the control experiment. (d) Wake state induction rate which indicates whether the wake state occurred within 100 s after light illumination. Upper column shows light phase and lower column shows dark phase. In each column, left side shows the activation group (blue light illumination) and right side shows the control group (yellow light illumination) (n = 3 mice, 5-10 manipulations/recordings, a total of 23 trials for activation [light phase], 18 trials for control [light phase], a total of 18 trials for activation [dark phase], 20 trials for control [dark phase]). (e) Wake latency after the initiation of optogenetic activation of VLS D1-MSNs during NREM sleep. The columns on the left and right show the light and dark phases, respectively. In each column, left bar shows the activation group, right bar shows the control group (n = 3 mice, 5-10 manipulations/recordings, a total of 23 trials for activation [light phase], 18 trials for control [light phase], a total of 23 trials for activation [dark phase], 30 trials for control [dark phase]; *p*_light_ = 0.01, *p*_dark_ = 0.03. independent *t*-test). (f) Percentage of wake time for 100 s after the initiation of VLS D1-MSNs photoactivation. The columns on the left and right show the light and dark phases, respectively. In each column, left bar shows the activation group, light bar shows the control group (n = 3 mice, 5-10 manipulations/recordings, a total of 23 trials for activation [light phase], 18 trials for control [light phase], a total of 23 trials for activation [dark phase], 30 trials for control [dark phase]; *p*_light_ = 6.3 × 10^-6^, *p*_dark_ = 2.4 × 10^-5^, independent *t*-test).

## Discussion

This study demonstrated that ablation of D2-MSN in the VLS causes an increase in the amount of wake time during the dark phase, accompanied by a decrease in sleep time. This effect becomes more pronounced as the ablation area expands to include the entire VS. Next, our fiber photometric recording revealed that the average intracellular Ca^2+^ signal level of the VLS D2-MSNs increased during the transition from wake to NREM sleep in mice and remained high during NREM and REM sleep compared to that in the wake state. Optogenetic activation of VLS D2-MSNs induced NREM sleep in mice from wake state, while VLS D1-MSN activation induced wake state from NREM sleep. These results suggest that D2-MSNs in the VLS have an NREM sleep-inducing function in coordination with those in other medial parts of the VS. Furthermore, this sleep-wake regulation function of VLS D2-MSNs was in the opposite direction to the D1-MSNs function in the same subregion.

Our ablation and optogenetic activation studies strongly suggest that D2-MSNs in the VLS have an NREM sleep-inducing function (**Figs. 2 and 5**), and the fiber photometric recording supports these results (**Fig. 4**). The VLS is distinguished from the medial parts of the VS, such as the NAc medial shell and core, based on cortical input patterns (Berendse et al., 1992; Hunnicutt et al., 2016). Several studies have demonstrated the functional specificities of D2-MSNs in the VLS, such as reward processing (Tsutsui-Kimura et al., 2017; Yang et al., 2018; Chen et al., 2021). However, our findings regarding the sleep-inducing function of D2-MSNs in the VLS are consistent with those found in the medial portion of the VS (NAc medial shell/core) (Satoh et al., 1999; Lazarus et al., 2011; Oishi et al., 2017; Luo et al., 2018).

Furthermore, we observed that the alteration of sleep/wake architecture was even more pronounced under the ablation of the D2-MSNs in the entire VS, compared to that of in the VLS (**Figs. 2a, b**). Therefore, the alterations in the sleep/wake pattern might be strengthened by the ablation of D2-MSNs in both the VLS and the medial part of the VS, indicating a coordinated and consistent regulation of the animal’s sleep-wake states by the D2-MSNs in these subregions of the VS. The degree of the D2-MSNs effect on the sleep/wake architecture may depend on the extent of its distribution area. D2-MSNs in the VS project to the VP (Basar et al., 2010), of which the projection from the NAc core to the VP has been reported to be involved in sleep-wake regulation (Oishi et al., 2017; Luo et al., 2018). These consistent efferent patterns could be associated with the common sleep-wake regulatory role of D2-MSNs in the medial and lateral VS.

We demonstrated that the optogenetic activation of the VLS D2-MSNs induced NREM sleep from wakefulness whereas that of the VLS D1-MSNs induced wake from NREM sleep in mice (**Figs. 5****, 6**). These findings strongly suggest that the VLS D1- and D2-MSNs activities themselves change the sleep-wake state of animals in the opposite direction. The reversal sleep-wake regulation function of the VLS D1- and D2-MSNs could be related to the differential dopaminergic control and/or output patterns of these two neuronal subpopulations. First, striatal D1- and D2-MSNs activities are inversely affected by the extracellular dopamine: As the D1 and D2 receptors couple with Gs and Gi proteins, respectively (Kebabian and Calne, 1979), the dopamine signal has excitatory and inhibitory effects on the activity of D1- and D2-MSNs, respectively.

Thus, dopaminergic control often causes the opposing function of striatal D1- and D2- MSNs, and their balance is expressed in functional outputs such as motor controls (Cox and Witten, 2019). It could be the same for the sleep-wake regulation function of the VLS D1- and D2-MSNs. Next, the partially differential output patterns of the VS D1- and D2-MSNs might cause their reversal of sleep-wake regulation functions. D2-MSNs in the VS consistently project to the VP, while the D1-MSNs in the NAc project to not only the VP but also the ventral mesencephalon (VM) (Kupchik et al., 2015; Liu et al., 2022). D1-MSNs in the NAc core have a wake-promoting role via the VM and lateral hypothalamus (Luo et al., 2018) and artificial inhibition of GABAergic neurons in the VM increases wakefulness time (Takata et al., 2018), whereas the VS D2-MSNs projection to the VP could involve NREM sleep induction (Oishi et al., 2017; Luo et al., 2018). These findings suggest that the VLS D2-MSNs activation could induce NREM sleep only through the VP whereas the VLS D1-MSNs activation-induced wake could be mediated by the inhibition of VM neurons.

We observed that the change in sleep/wake architecture caused by VLS/VS D2- MSNs ablation occurred only during the dark phase (**Fig. 2**). Previous studies also reported that the inhibition D2-MSNs in the NAc core induced wake state in mice during the dark phase, although they did not mention during the light phase (Oishi et al., 2017; Luo et al., 2018). The sleep-inducing function of D2-MSNs restricted to the dark phase may be associated with striatal dopaminergic signaling. The extracellular dopamine levels in the striatum exhibit a diurnal variation, with lower levels during the light phase and higher levels during the dark phase (Hood et al., 2010). Striatal extracellular dopamine reversely affects the D2- and D1-MSNs activities, and the activity/function of D1-MSNs relative to that of D2-MSNs is strengthened by increased extracellular dopamine during the dark phase, which might be further augmented by the D2-MSNs ablation, causing a dark phase-specific increase in wakefulness. Besides, we observed that optogenetic activation of VLS D2- and D1-MSNs promptly induced sleep-wake state change regardless of the light or dark phases (**Figs. 5****, 6**). These findings suggest that the firing activity of D1/D2-MSNs may be capable of inducing changes in the sleep/wake states, independent of the light/dark phases.

Our fiber photometric recordings revealed that the compound intracellular Ca^2+^ signal in the VLS D2-MSNs was higher on average during NREM and REM sleep than in the wake state, and the mean signal level was elevated during the wake-to-NREM sleep transition (**Fig. 4**). These state-dependent Ca^2+^ signal variations in the VLS D2- MSNs strongly support our conclusion of the sleep-inducing function of the VLS D2- MSNs derived from our ablation and optogenetic experiments (**Figs. 2 and 5**). The VLS D2-MSNs activity can be affected by dopaminergic input from the ventral tegmental area (VTA; Mingote et al., 2019) and glutamatergic input from the insular cortex (Berendse et al., 1992). The activity of VTA dopaminergic neurons varies depending on the sleep-wake state, which is higher during wakefulness and REM sleep and lower during NREM sleep (Eban-Rothschild et al., 2016). Therefore, VTA dopaminergic inputs, which could suppress D2-MSNs activity via the Gi-coupled D2 receptor, could cause the VLS D2-MSNs activity patterns to be lower during wakefulness and higher during NREM sleep. However, the higher activity of VLS D2-MSNs during REM sleep cannot be explained by variations in the dopaminergic input. The increase in the Ca^2+^ signal level of the VLS D2-MSNs during REM sleep may be influenced by glutamatergic neuronal input from the insular cortex. A previous study showed that lesions in the insular cortex led to decreased wake time and increased NREM and REM sleep times (Chen et al., 2016). The fluctuation of neuronal activity in the insular cortex across sleep-wake states is currently unclear; however, given its potential involvement in sleep/wake regulation, it is not unexpected that glutamatergic neuronal projections from the insular cortex may impact the activity of D2-MSNs in the VLS across different sleep-wake states. In addition, these diverse state-dependent neuronal inputs might cause detailed differences in the VLS D2-MSNs Ca^2+^ signal fluctuations between each sleep-wake state in terms of their relationship with EEG/EMG activity in mice (**Figs. 4e, f**). Further studies are required to elucidate the sleep-inducing functions of the VLS D2- MSNs at the neuronal circuit level.

Our study has two limitations. First, the D2-MSN ablation method in D2-DTA mice (Tsutsui-Kimura et al., 2017) cannot evaluate the sleep-wake regulatory function of D2-MSNs in each subregion of the VS separately, except for the VLS. Several prior studies have provided data on D2-MSNs in various subregions of the VS that were not included in our study (Oishi et al., 2017; Luo et al., 2018). Next, it has been pointed out that fluorescent probes expressed in the living brain and detected by the fiber photometry system could be affected by cerebral blood volume (Ikoma et al., 2023a, 2023b). The evaluation of YC signal fluctuations particularly during REM sleep should be cautious because cerebral blood flow is increased (Braun et al., 1997; Tsai et al., 2021; Ikoma et al., 2023b). The proposed new fiber photometry setup and analysis method to exclude the effects of cerebral blood flow (Ikoma et al., 2023a, 2023b) could improve the accuracy of our neuronal activity assessment based on the YC signal measurement in the future.

In conclusion, our results demonstrate that D2-MSNs in the VLS are essential for the maintenance of sleep/wake architecture and have an NREM sleep-inducing function.

## Acknowledgments

This work was supported by the Japan Science and Technology Agency (JST) Strategic Basic Research Program (CREST) “Research on Multi-sensing Biosystems and Development of Adaptive Technologies” (22gm1510007h0001) from AMED to KFT. We would like to express our gratitude to Professor Yasue Mitsukura for providing valuable technical advice and engaging discussions throughout this study. We also extend our appreciation to Mr. Kaiki Matsuo for proofreading the manuscript.

## Abbreviations

VLS D2-MSNs: control sleep-wake state

## References

1. Basar K, Sesia T, Groenewegen H, Steinbusch HWM, Visser-Vandewalle V, Temel Y (2010) Nucleus accumbens and impulsivity. Prog Neurobiol 92:533–557.

2. Berendse HW, Galis-de Graaf Y, Groenewegen HJ (1992) Topographical organization and relationship with ventral striatal compartments of prefrontal corticostriatal projections in the rat. J Comp Neurol 316:314–347.

3. Berndt A, Yizhar O, Gunaydin LA, Hegemann P, Deisseroth K (2009) Bi-stable neural state switches. Nat Neurosci 12:229–234.

4. Braun AR, Balkin TJ, Wesenten NJ, Carson RE, Varga M, Baldwin P, Selbie S, Belenky G, Herscovitch P (1997) Regional cerebral blood flow throughout the sleep- wake cycle. Brain. 120:1173–1197.

5. Chen R, Blosser TR, Djekidel MN, Hao J, Bhattacherjee A, Chen W, Tuesta LM, Zhuang X, Zhang Y (2021) Decoding molecular and cellular heterogeneity of mouse nucleus accumbens. Nat Neurosci 24:1757–1771.

6. Chen M, Chiang WY, Yugay T, Patxot M, Özcivit IB, Hu K, Lu J (2016) Anterior Insula Regulates Multiscale Temporal Organization of Sleep and Wake Activity. J Biol Rhythms 31:182–193.

7. Choi JH, Koch KP, Poppendieck W, Lee M, Shin H-S (2010) High resolution electroencephalography in freely moving mice. J Neurophysiol 104:1825–1834.

8. Cox J, Witten IB (2019) Striatal circuits for reward learning and decision-making. Nat Rev. Neurosci 20:482–494.

9. Durieux PF, Bearzatto B, Guiducci S, Buch T, Waisman A, Zoli M, Schiffmann SN, de Kerchove d’Exaerde A (2009) D2R striatopallidal neurons inhibit both locomotor and drug reward processe. Nat Neurosci 12:393–395.

10. Eban-Rothschild A, Rothschild G, Giardino WJ, Jones JR, Lecea L (2016) VTA dopaminergic neurons regulate ethologically relevant sleep–wake behaviors. Nat Neurosci 19:1356–1366.

11. Funato H, Miyoshi C, Fujiyama T, Kanda T, Sato M, Wang Z, Ma J, Nakane S, Tomita J, Ikkyu A, Kakizaki M, Hotta-Hirashima N, Kanno S, Komiya H, Asano F, Honda T, Kim SJ, Harano K, Muramoto H, Yonezawa T, Mizuno S, Miyazaki S, Connor L, Kumar V, Miura I, Suzuki T, Watanabe A, Abe M, Sugiyama F, Takahashi S, Sakimura K, Hayashi Y, Liu Q, Kume K, Wakana S, Takahashi JS, Yanagisawa M (2016) Forward-genetics analysis of sleep in randomly mutagenized mice. Nature 539:378–383.

12. Hood S, Cassidy P, Cossette MP, Weigl Y, Verwey M, Robinson B, Stewart J, Amir S (2010) Endogenous Dopamine Regulates the Rhythm of Expression of the Clock Protein PER2 in the Rat Dorsal Striatum via Daily Activation of D2 Dopamine Receptors. J Neurosci 30:14046–14058.

13. Horikawa K, Yamada Y, Matsuda T, Kobayashi K, Hashimoto M, Matsuura T, Miyawaki A, Michikawa T, Mikoshiba K, Nagai T (2010) Spontaneous network activity visualized by ultrasensitive Ca^2+^ indicators, yellow Cameleon-Nano. Nat Methods 7:729–732.

14. Huang ZL, Zhang Z, Qu WM (2014) Roles of adenosine and its receptors in sleep-wake regulation. Int Rev Neurobiol 119:349–371.

15. Hunnicutt BJ, Jongbloets BC, Birdsong WT, Gertz KJ, Zhong H, Mao T (2016) A comprehensive excitatory input map of the striatum reveals novel functional organization. eLife 5:e19103.

16. Ikoma Y, Sasaki D, Matsui K (2023a) Local brain environment changes associated with epileptogenesis. Brain. 146 (2): 576–586.

17. Ikoma Y, Takahashi Y, Sasaki D, Matsui K (2023b) Properties of REM sleep alterations with epilepsy. Brain. awac499.

18. Kanemaru K, Sekiya H, Xu M, Satoh K, Kitajima N, Yoshida K, Okubo Y, Sasaki T, Moritoh S, Hasuwa H, Mimura M, Horikawa K, Matsui K, Nagai T, Iino M, Tanaka KF (2014) In vivo visualization of subtle, transient, and local activity of astrocytes using an ultrasensitive Ca^2+^ indicator. Cell Rep 8:311–318.

19. Kato T, Mitsukura Y, Yoshida K, Mimura M, Takata N, Tanaka KF (2022) Oscillatory Population-Level Activity of Dorsal Raphe Serotonergic Neurons Is Inscribed in Sleep Structure. J Neurosci 42:7244–7255.

20. Kebabian JW, Calne DB (1979) Multiple receptors for dopamine. Nature 277:93–96.

21. Kemp JM, Powell TP (1971) The structure of the caudate nucleus of the cat: light and electron microscopy. Philos Trans R Soc Lond B Biol Sci 262:383–401.

22. Kravitz AV, Freeze BS, Parker PR, Kay K, Thwin MT, Deisseroth K, Kreitzer AC (2010) Regulation of parkinsonian motor behaviours by optogenetic control of basal ganglia circuitry. Nature 466:622–626.

23. Kupchik YM, Brown RM, Heinsbroek JA, Lobo MK, Schwartz DJ, Kalivas PW (2015) Coding the direct/indirect pathways by D1 and D2 receptors is not valid for accumbens projections. Nat Neurosci 18:1230–1232.

24. Lazarus M, Shen H-Y, Cherasse Y, Qu W-M, Huang Z-L, Bass CE, Winsky-Sommerer R, Semba K, Fredholm BB, Boison D, Hayaishi O, Urade Y, Chen JF (2011) Arousal effect of caffeine depends on adenosine A2A receptors in the shell of the nucleus accumbens. J Neurosci 31:10067–10075.

25. Lee P, Morley G, Huang Q, Fischer A, Seiler S, Horner JW, Factor S, Vaidya D, Jalife J, Fishman GI (1998) Conditional lineage ablation to model human diseases. PNAS 95:11371–11376.

26. Liu Z, Le Q, Lv Y, Chen X, Cui J, Zhou Y, Cheng D, Ma C, Su X, Xiao L, Yang R, Zhang J, Ma L, Liu X (2022) A distinct D1-MSN subpopulation down-regulates dopamine to promote negative emotional state. Cell Res 32(2): 139–156.

27. Luo Y-J, Li Y-D, Wang L, Yang S-R, Yuan X-S, Wang J, Cherasse Y, Lazarus M, Chen J-F, Qu W-M, Huang Z-L (2018) Nucleus accumbens controls wakefulness by a subpopulation of neurons expressing dopamine D1 receptors. Nat Commun 9:1576.

28. McCoy EJ, Walden AT (1998) Multitaper Spectral Estimation of Power Law Process. IEEE Transactions on Signal Processing. 46:655–658.

29. Mingote S, Amsellem A, Kempf A, Rayport S, Chuhma N (2019) Dopamine-glutamate neuron projections to the nucleus accumbens medial shell and behavioral switching. Neurochem Int 129:104482.

30. Natsubori A, Tsutsui-Kimura I, Nishida H, Bouchekioua Y, Sekiya H, Uchigashima M, Watanabe M, de Kerchove d’Exaerde A, Mimura M, Takata N, Tanaka KF (2017) Ventrolateral striatal medium spiny neurons positively regulate food-incentive, goal- directed behavior independently of D1 and D2 selectivity. J Neurosci 37:2723–2733.

31. Oishi Y, Lazarus M (2017) The control of sleep and wakefulness by mesolimbic dopamine systems. Neurosci Res 118:66–73.

32. Oishi Y, Xu Q, Wang L, Zhang B-J, Takahashi K, Takata Y, Luo Y-J, Cherasse Y, Schiffmann SN, de Kerchove d’Exaerde A, Urade Y, Qu W-M, Huang Z-L, Lazarus M (2017) Slow-wave sleep is controlled by a subset of nucleus accumbens core neurons in mice. Nat Commun 8:734.

33. Satoh S, Matsumura H, Koike N, Tokunaga Y, Maeda T, Hayaishi O (1999) Region- dependent difference in the sleep-promoting potency of an adenosine A2A receptor agonist. Eur J Neurosci 11:1587–1597.

34. Sano H, Nagai Y, Miyakawa T, Shigemoto R, Yokoi M (2008) Increased social interaction in mice deficient of the striatal medium spiny neuron-specific phosphodiesterase 10A2. J Neurochem 105:546–556.

35. Takata Y, Oishi Y, Zhou XZ, Hasegawa E, Takahashi K, Cherasse Y, Sakurai T, Lazarus M (2018) Sleep and Wakefulness Are Controlled by Ventral Medial Midbrain/Pons GABAergic Neurons in Mice. J Neurosci. 38(47): 10080–10092.

36. Tsai CJ, Nagata T, Liu CY, Vogt KE, Yanagisawa M, Hayashi Y (2021) Cerebral capillary blood flow upsurge during REM sleep is mediated by A2a receptors. Cell Reports 36:109558.

37. Tanaka KF, Samuels BA, Hen R (2012) Serotonin receptor expression along the dorsal– ventral axis of mouse hippocampus. Philos Trans R Soc Lond B Biol Sci 367:2395– 2401.

38. Tobler I, Deboer T, Fischer M (1997) Sleep and Sleep Regulation in Normal and Prion Protein-Deficient Mice. J Neurosci 17:1869–1879.

39. Tsutsui-Kimura I, Natsubori A, Mori M, Kobayashi K, Drew MR, d’Exaerde AK, Mimura M, Tanaka KF (2017) Distinct Roles of Ventromedial versus Ventrolateral Striatal Medium Spiny Neurons in Reward-Oriented Behavior. Curr Biol 27:3042–3048.

40. Tsutsui-Kimura I, Takiue H, Yoshida K, Xu M, Yano R, Ohta H, Nishida H, Bouchekioua Y, Okano H, Uchigashima M, Watanabe M, Takata N, Drew MR, Sano H, Mimura M, Tanaka KF (2017) Dysfunction of ventrolateral striatal dopamine receptor type 2-expressing medium spiny neurons impairs instrumental motivation. Nat Commun 8:14304.

41. Yacoubi ME, Ledent C, Menard JF, Parmentier M, Costentin J, Vaugeois JM (2000) The stimulant effects of caffeine on locomotor behaviour in mice are mediated through its blockade of adenosine A2A receptors. Br J Pharmacol 129:1465–1473.

42. Yang H, Jong JW, Tak Y, Peck J, Bateup HS, Lammel S (2018) Nucleus Accumbens Subnuclei Regulate Motivated Behavior via Direct Inhibition and Disinhibition of VTA Dopamine Subpopulations. Neuron 97:434–449.

43. Yoshida K, Tsutui-Kimura I, Kono A, Yamanaka A, Kobayashi K, Watanabe M, Mimura M, Tanaka KF (2020) Opposing Ventral Striatal Medium Spiny Neuron Activities Shaped by Striatal Parvalbumin-Expressing Interneurons during Goal- Directed Behaviors. Cell Reports 31:107829.

44. Yuan XS, Wang L, Dong H, Qu W-M, Yang S-R, Cherasse Y, Lazarus M, Schiffmann SN, de Kerchove d’Exaerde A, Li R-X, Huang Z-L (2017) Striatal adenosine A2A receptor neurons control active-period sleep via parvalbumin neurons in external globus pallidus. eLife 6:e29055.

